# Grove-level analysis of titer and prevalence of “*Candidatus* Liberibacter asiaticus” and *Wolbachia* in *Diaphorina citri,* vector of citrus Huanglongbing

**DOI:** 10.1101/2023.10.04.560920

**Authors:** Marina Mann, Luke Thompson, Lynn Johnson, Michelle Heck

## Abstract

Huanglongbing (HLB, or citrus greening disease) affects all citrus varieties world-wide. In the USA, Asia, and South America the causal agent is “*Candidatus* Liberibacter asiaticus” (*C*Las), a phloem-limited, uncultured, alphaproteobacterium. The hemipteran insect vector, *Diaphorina citri* (Asian citrus psyllid) acquires and transmits *C*Las in a circulative, propagative manner. In addition to *C*Las, *D. citri* hosts multiple symbiotic bacterial species including *Wolbachia* (wDi). In *D. citri*, wDi has been sequenced and studied but specific roles in *D. citri* biology are unknown. Using well established quantitative PCR methods we measured *C*Las titer in *D. citri* collected from four groves in central Florida with distinct HLB management strategies and tested whether *C*Las and wDi titer were correlated in a sub-set of these insects. Grove site had the largest effect on *C*Las titer. Sex had no effect on *C*Las titer, while higher wDi titer was correlated with non-infected insects. Our results suggest that more directed follow-up research is necessary and important to clarify whether field management tactics influence *C*Las titer in *D. citri* and to better understand gene-by-environment interactions among *D. citri*, wDi and *C*Las. Now that millions of trees in Florida have been treated with injectable formulations of oxytetracycline, which is likely to decrease bacterial populations in *D. citri*, this study may represent the last biologically meaningful snapshot of grove-level vector-pathogen ecology in the state during the HLB epidemic.

## Introduction

Huanglongbing (HLB, also known as citrus greening disease) affects all citrus varieties world-wide, a total of 61 countries as of December 2022) (Ghosh et al. 2023) and after decades of targeted research for solutions and over $1 billion US dollars spent in research, HLB remains the most devastating disease of citrus with no cure. It is caused by an unculturable, intracellular, phloem-limited bacteria in the genus “*Candidatus* Liberibacter”. The predominant species causing HLB is “*Ca.* L. asiaticus” (*C*Las). Two other species, “*Ca.* L. africanus” and “*Ca.* L. americanus” have also been described in multiple African countries and South-Central American countries, respectively. *C*Las is spread from tree to tree by the insect vector *Diaphorina citri* (Asian citrus psyllid), a hemipteran in the family Psyllidae which feeds exclusively on the phloem of citrus trees as well as species of *Murraya*. *C*Las is transmitted by *D. citri* in an intracellular, circulative, propagative manner (Ammar et al. 2016; Inoue et al. 2009; Higgins et al. 2022). While *C*Las can also spread via grafting infected citrus budwood, vector-borne dispersal is the predominant and least controllable method causing new citrus infections. Lastly, *C*Las can infect periwinkle via dodder, but periwinkle is not a host for *D. citri* (Garnier and Bove 1983).

Management of the vector requires an understanding of basic epidemiology and vector biology in the field. In the United States, strict testing regimens are followed in Texas and California to maintain quarantine areas for the vector and pathogen. But HLB is now endemic in Florida. Despite the frequency of psyllid testing and a known increase in insecticide resistance by *D. citri* (Chen et al. 2022; Vázquez-García et al. 2013; Naeem et al. 2019; Rao et al. 2014; Pardo et al. 2018; Venkatesan et al. 2022), relatively few papers have been published on population-level dynamics of *C*Las-*D. citri* infections in the field. A few studies have looked at *D. citri-C*Las titer and prevalence in different situations, including specific field sites, month to month changes, year to year trends, and differences between countries (Bové et al. 1993; Wulff et al. 2020; Capoor et al. 1974; Coy and Stelinski 2015; Hall 2018; Manjunath et al. 2008; Razi et al. 2014; Tiwari et al. 2010; Sétamou et al. 2020; Ukuda-Hosokawa et al. 2015; Walter et al. 2012). These studies reported high levels of variation in *C*Las titer across both the individual and population level. Genetic studies have also shown that psyllid populations vary in their *C*Las acquisition, transmission and titer phenotypes and that this variation is stable and heritable over many generations, consistent with these phenotypes being regulated by *D. citri* genes and proteins (Ramsey et al. 2015; Mann et al. 2018; Ammar et al. 2018).

In addition to *C*Las, *D. citri* hosts multiple symbiotic bacteria which it maintains because they provide various benefits (such as nutritional supplements and defensive polyketides) (Hosseinzadeh et al. 2019b; Nakabachi et al. 2013). *Wolbachia* (wDi) is one such endosymbiont found in *D. citri*, as well as over 50% of all insects. At least two genetically distinct populations of wDi have been shown to infect *D. citri* in Florida (Chu et al. 2019). Understanding how wDi interacts with both *D. citri* and *C*Las may be important to the future management of psyllid population (Chu et al. 2019; Jain et al. 2017). In *D. citri*, wDi has been sequenced (Neupane et al. 2022; Saha et al. 2012; Shippy et al. 2019) and studied but its specific role in *D. citri* is still unclear. Studies suggest wDi is 100% present in *D. citri* individuals (Hosseinzadeh et al. 2019b; Guidolin and Cônsoli 2013; Hoffmann et al. 2014) and is found infecting many organs (Hosseinzadeh et al. 2019b; Ren et al. 2018) of *D. citri*. The bacteriome, the endosymbiotic organ where the other psyllid endosymbiont bacteria reside, maintains lower titers as compared to other organs (Hosseinzadeh et al. 2019b).

The goal of this study was to compare *C*Las titer of psyllids collected from four groves in Florida in 2018 with high disease pressure to better understand the natural variation that exists in *C*Las titer across different populations of *D. citri* and to test whether factors such as sex or wDi titer influence *C*Las titer.

## Methods

### Sample collection

Four citrus groves in central Florida, near lake Okeechobee, were sampled for adult *Diaphorina citri* during the first week of October, 2018 (see **Figure S1**). Individual *D. citri* were collected into 70% ethanol using mouth aspiration. At least 1200 individual *D. citri* were collected from each field. Samples were air dried and randomly aliquoted into groups of 100, promptly flash frozen in liquid nitrogen and stored at -80 *℃* until DNA extractions.

### DNA extraction and quantification

DNA extractions were performed in three batches on a total of 1062 samples, randomly selected from each field site in varying amounts, using the Qiagen DNeasy Blood and Tissue 96 extraction kit. This kit allowed 96 samples to be extracted simultaneously with a filter-based method. The methods advertised with the kit were followed with minor adjustments as follows. First, individual *D. citri* were sexed under a dissecting scope over ice, then moved via paint brush to the 96-well racks of sample tubes supplied by the Qiagen kit. Three, 3.2mm steel beads were added to each tube and the samples were capped using the rubber tube caps provided, throughout the first and last steps of the protocol. Samples were disrupted for 2 minutes at 1000 rpm using a mechanical homogenization with a SPEX Sample Prep MiniG 1600 (Thermo Fisher). The samples were kept on ice before and after sample disruption. All centrifugation steps were performed at the maximum 4000 rpm (RCF of 3434 x g) available when using the Hermle Z466 3L swing-out rotor with square microplate buckets and inserts to accommodate the stacked column and flow-through plates. Samples were lysed for 45 min at 56 *℃* in a water bath and mixed once midway. Following lysis, 4 ul RNase A per sample was added to purify the genomic DNA, at which point, instead of tube caps, plastic PCR plate covers were used to cover the 96-well racks. Samples were eluted once using 50 ul AE buffer then stored at -20 *℃* until DNA quantification and purity checks, which were performed using a Nanodrop 2000.

### Quantitative PCR

Quantitative polymerase chain reaction (qPCR) was performed on all 1062 samples using TaqMan hydrolysis probes, testing for the presence and relative titer quantification of the “*Candidatus* Liberibacter asiaticus” (*C*Las) 16S rRNA gene. The 16S rRNA forward primer (51-TCGAGCGCGTATGCAATACG-31), reverse primer (51-GCGTTATCCCGTAGAAAAAGGTAG-31) and probe (51-FAM-AGACGGGTG/ZEN/AGTAACGCG-31) were used, as previously established (Wenbin Li, 2006). The probe includes a fluorescent dye (51 6-FAM) and a quencher (ZEN, 31 Iowa Black FQ). Standard thermal conditions were used during the qPCR. Specifically, 95 °C for 10 min (1 cycle) and 95 °C for 15 sec and 60 °C for 60 sec (40 cycles total). Each qPCR 384-well plate included a standard curve dilution series (a serial dilution from 10^9^ down to 10^3^ of the synthetic plasmid pUC57 containing a *C*Las 16S rDNA gene fragment). The resulting y-intercept was 38.303, R^2^ = 0.964, slope = -3.095, and efficiency = 110.452%. Each qPCR plate also contained positive and negative controls run with two technical replicates each. A subset of samples from the first batch were subjected to additional qPCR using SYBR Green reagents to access wDi presence-absence and estimate titer. The wDi *ftsZ* gene (ftsZ-81) (A. S. Guidolin 2013) forward primer (5’AGCAGCCAGAGAAGCAAGAG3’) and reverse primer (5’TACGTCGCACACCTTCAAAA3’), and the *Wingless* (*WgL*) (MyLo L. Thao 2000) forward primer (GCTCTCAAAGATCGGTTTGACGG) and reverse (GCTGCCACGAACGTTACCTTC) primers were run in parallel on the same 386 well plate. All samples on every qPCR plate were run in duplicate (2 technical replicates) then averaged to create the reported final quantitative cycle (Cq) values. The maximum number of qPCR cycles is 40, where Cq = 40 means the primers did not create a product that could be amplified so the sample is non-infected or titers are below the limit of detection, while Cq < 40 suggests copies of the target gene are present, and the insect is infected. *C*Las cell counts were calculated from Cq using the standard curve and the QuantStudio6 software. Each qPCR well contained 20 ng of DNA from a single *D. citri.* The QuantStudio6 software associated with the qPCR machine used the standard curve dilution values and resulting best-fit linear equation and efficiency metrics to internally calculate the *C*Las cell equivalent to the number of copies of the *C*Las rDNA sequence in a single *D. citri*. The values were exported from the software as an excel document. *C*Las cell equivalents for technical replicates of each sample were averaged (note, Cq = 40 yielded undetermined values, which were transformed to zeros for calculations of cell equivalents), then all samples from each field were averaged. Field mean *C*Las cell equivalent was found by taking the difference.

### Statistical Methods

A linear mixed-effects model (LMM) was used to analyze the *C*Las Cq data collected on the individual insects… Two models were compared to determine whether grove makes a significant difference as a random effect. Model 1 included both DNA extraction batch and grove as random effects, and sex as a fixed effect. Model 2 was reduced, containing only batch as a random effect and sex as a fixed effect, with grove absent. An ANOVA was used to test the difference between the two models. Model assumptions of normality and homogeneous variance were assessed visually using the model residuals. *P*-values < 0.05 were considered statistically significant. The mean absolute titer of 918 individuals was calculated from a standard curve, then the mean absolute titer was calculated for each field site. For each pairwise field comparison, non-zero titers were used to calculate field-specific means, the means subtracted to find the difference and the absolute value was reported to highlight the magnitude of the difference the second from the first (**Figure 2**). A subset of the samples from batch number one, representing all four fields, was additionally tested for the dual effect of wDi (Cq values normalized by *Wingless* Cq) and grove on *C*Las Cq. R packages (R Core Team 2022) were required for the analysis of both *C*Las and wDi, including *ggplot2* (Wickham 2-16)*, ggpubr* (Kassambara 2020), and *lmertest* (Kuznetsova et al. 2017). To test for interactions between *C*Las and wDi in the insects, wDi Cq and *C*Las Cq were converted into binary values as follows: for wDi, a value of 0 was assigned for Cq values below the median, and a value of 1 was assigned for values above median and for *C*Las, a value of 0 was assigned for Cq values less than 40, and a value of 1 for undermined Cq values or Cq=40). The association between wDi and *C*Las titer was assessed using a Wilcoxon test, which clarified whether wDi Cq above and below the median was different in CLas (+) and (-) insects, and a Chi-squared test in R to test whether the wDi Cq distribution around the median differs from the distributions of *C*Las Cq below or at Cq=40.

## Results

### 1. Each grove had a distinct characteristics, including tree age, type and management strategies

Groves were assigned names corresponding to their town of origin to protect the identity of the participating growers. Psyllid infestations were found by scouting for flush points with nymph honeydew deposits. A total of 450 adult *D. citri* were used from the Port St. Lucie grove. This grove was one of two organically managed groves we sampled from, however, Port St. Lucie did not utilize any pesticides, herbicides or other chemical inputs to our knowledge, at the time of sampling. It maintained a high *D. citri* population as well as the highest observed levels of in-field entomopathogenic fungi (Higgins et al. 2022) and other insects (including ladybeetles, ants, caterpillars and wasps). Multiple citrus species as well as *Murraya spp.* grew within the grove. The majority of *D. citri* samples were taken from *Citrus sinensis* (Valencia orange). Trees ranged in age from recently re-planted to well established, bearing trees. We used a total of 164 *D. citri* adults from *C. sinensis* in the second grove site, referred to as Pioneer. This grove was also organically managed, and while organic pesticides were used, only a regimen of periodic nitrogen, phosphorous and potassium (NPK) supplementation was described in particular during sample collection, and any other treatments utilized are unknown. Trees were mature and bearing and well-spaced out. A total of 176 samples were used from an experimental plot within a third grove, Whidden Corner. This grove was composed of mostly small, juvenile, recently planted *C. sinensis* trees undergoing heavy pesticide treatment for *D. citri* control, with applications occurring every three weeks. A total of 272 adult *D. citri* were used from a fourth citrus grove, referred to as Montura. These insects were sampled from *C. sinensis,* and this grove also experienced insecticide sprays every three weeks. The trees were bearing, tall, and thick with leaves. In all groves, rootstock type was unknown, and the majority of *D. citri* were collected from flush.

### 2. Grove had the largest effect on *C*Las Cq

The random effect of grove site significantly outweighed any effect of sex or DNA extraction batch on *C*Las Cq (Chi^2^ = 462.01, df = 2, *P* < 0.001). The effect of DNA extraction batch was included in the model to rule out a potential bias, and we found no between-batch variability (var. = 0.010, SD = 0.010) suggesting high accuracy of technical replications of DNA extraction.

The four populations of *D. citri* fell into three broad categories based on *C*Las titer and secondarily, infection rates. After accounting for effects of batch and sex, Pioneer represents the “low titer” population and has the lowest overall titer (bacterial load per individual *D. citri*) with mean Cq = 39.0 and standard error (SE) = 0.32. Montura has the highest titer (mean Cq = 30.4, SE = 0.28) and represents the “high titer” population, while Port St. Lucie (mean Cq = 32.1, SE = 0.30) and Whidden Corner (mean Cq = 31.6, SE = 0.30) report similar bacterial titers and make up the third population of “medium titer” individuals. When absolute titer of *C*Las is calculated from Cq and a standard curve, the three groups are still present. Means were calculated using only samples with Cq<40, and Montura had the highest titer (29,630 *C*Las cell equivalents), Pioneer the lowest titer (41 *C*Las cell equivalents), while Whidden Corner and Port St. Lucie fell in the middle (8,912 and 11,270 *C*Las cell equivalents, respectively).

The percent infection rate is calculated using the proportion of insects with Cq values lower than 40, as Cq = 40 are considered non-infected with *C*Las. While the three groups above are determined by *C*Las titer, their infection rates do not necessarily match the trend. The “low titer” population represented by Pioneer did also have the lowest *C*Las infection rate (26.82 %). However, the “medium titer” populations represented by Port St. Lucie (97.11 %) and Whidden Corner (96.02 %) recorded the highest percent infection rates. Finally, the “high titer” population represented by Montura maintained the second-lowest percent infection rate (94.85 %). The three populations as determined by mean Cq can be seen in **Figure 1** and pair-wise results between grove populations (**Figure 2**) additionally confirm visual differences between groves.

**Figure 1.**
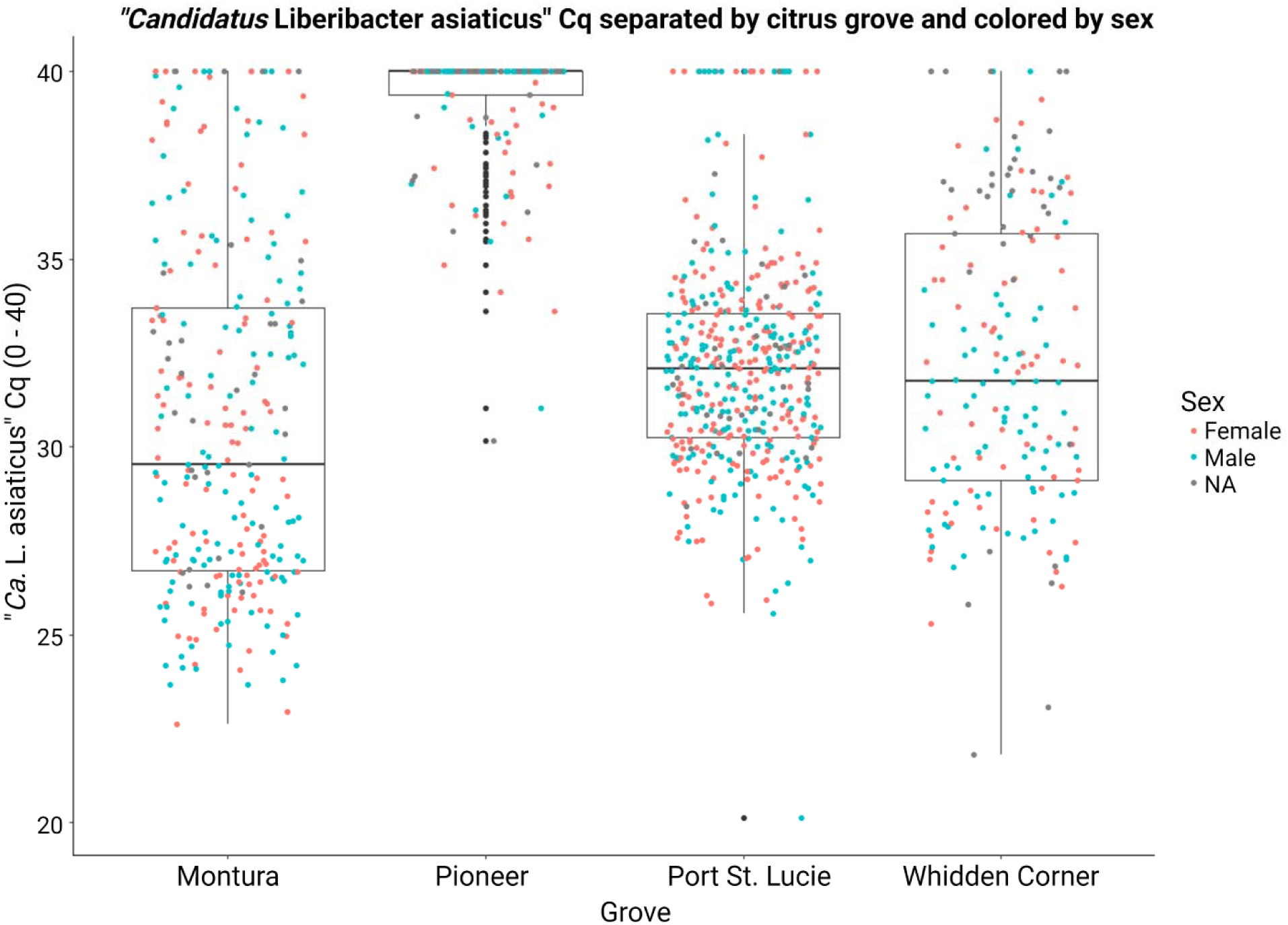
Box plot of CLas Cq distribution by citrus grove collection site and colored by sex. Box plot center lines are the median, box top and bottom edges are the first and third quartile respectively, and outliers are points that extend past the tails. NA samples (grey) are those that have a CLas Cq value but no recorded sex. M = male (blue), F = female (red).

**Figure 2:**
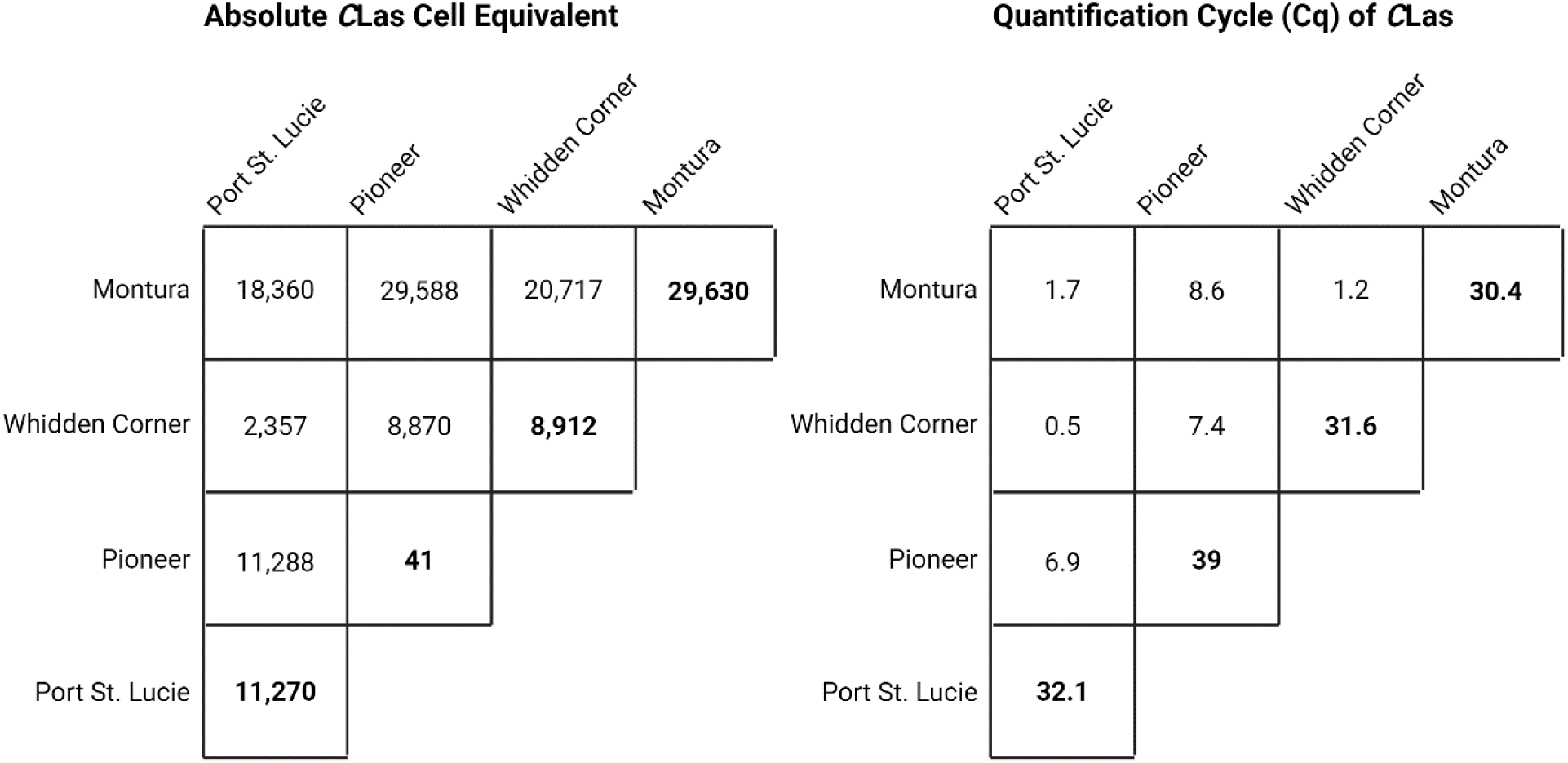
Pair-wise field comparisons using both mean CLas Cq and titer of CLas (genome copies of CLas in 20 ng of *D. citri* DNA). Field-specific means were computed using only non-zero absolute titer values, or Cq < 40 values and the absolute values are reported.

### 3. The sex of *D. citri* and color morph had no effect on CLas Cq means

Sex of each *D. citri* individual was recorded. Of the 929 samples with recorded sex, the overall ratio of males to females was nearly equal (Male: n=454, 48.86%; Female: n=475, 51.13%) as was the ratio of males to females in each grove (Montura: Female=41.54%, n=113, Male=46.32%, n=126; Pioneer: Female=42.68%, n=70, Male=39.02%, n=64; Port St. Lucie: Female=50.00%, n=225, Male=42.22%, n=190; Whidden Corner: Female=38.07%, n=67, Male=42.05%, n=74) (see **Table S1A** for further breakdown of male-female results).

The effect of sex on CLas Cq was tested and included in our model as CLas infection ha been shown to affect *D. citri* behavior differently by sex. There was no effect of sex on CLas Cq after controlling for grove and DNA extraction batch (F = 0.489, df =1, *P* = 0.484). This can also be seen in **Figure 1**, as the blue and red colors show a random distribution of sex by Cq value for each of the four groves.

In laboratory experiments, the color morph of *D. citri* was shown to be associated with *C*Las titers (Hosseinzadeh et al. 2019a). While it was technically challenging to accurately assign the color morphology to each specimen due to the insects being stored in ethanol prior to data collection, no obvious associations between *C*Las titer and color morph emerged in these insects (**Table S1B**). The lack of association between color morphology and *C*Las infection status has previously been reported in the analysis of other grove-captured, Florida psyllid populations (Hall et al. 2018).

### 4. *Wolbachia* (wDi) has higher titer in non-infected *D. citri*

A subset of *D. citri* samples from the initial DNA extraction were tested for wDi titer. The total number of samples tested was 176 (Port St, Lucie = 46, Pioneer = 42, Montura = 41, Whidden Corner = 47). A linear regression compared wDi Cq to *C*Las Cq, revealing no correlation between *C*Las and wDi Cq among *D. citri* from these Florida citrus groves (R^2^ = 0.0005) (**Figure S2**). Additionally, wDi Cq was normalized against the *D. citri Wingless* gene (**Figure S3**). A linear model assessed the relative effects of both grove and normalized wDi Cq on *C*Las Cq. The analysis indicated that wDi was not significantly correlated with *C*Las Cq (F = 0.646, df = 1, *P* = 0.426). Moreover, the source grove had the most substantial effect on *C*Las titer (F = 22.881, df = 3, *P* < 0.001, **Table S2**).

During the regression analysis, many wDi samples exhibited low Cq values when *C*Las Cq = 40 (indicating non-infected samples, see **Figure S2**). We investigated if wDi titer differed significantly based on *C*Las presence (defined as Cq < 40) or absence (Cq = 40 Our analysis revealed that when *C*Las is absent, wDi Cq was more likely to be below the median wDi Cq value (Wilcoxon, n=119, *P* = 0.017). Furthermore, a Chi^2^ test confirmed a significant difference in the proportion of wDi samples with Cq above and below the median *C*Las Cq distribution, when that distribution is divided by infection or no infection (X^2^(1, n = 119) = 6.086, *P* = 0.013). This exploratory testing appears to show that non-infected *D. citri* tend to have higher wDi titers compared to *C*Las-infected individuals.

## Discussion

Natural variation in vector competency plays a role in disease epidemiology. Vector competency is the potential of a vector to transmit a pathogen. In *D. citri*, vector competency is in part regulated by *D. citri* genes and proteins functioning at distinct tissue barriers which *C*Las must overcome prior to inoculation into a new host tree (Ammar et al. 2018; Mann et al. 2018; Kruse et al. 2017, 2018; Hosseinzadeh and Heck 2023). The infection of the host tree and citrus variety has also been shown to play a role in modulating vector capacity (Ramsey et al. 2022; Chin et al. 2020; Lombardi et al. 2024; Coletta-Filho et al. 2014). *D. citri* acquires and transmits *C*Las in an intracellular, circulative, propagative manner. Vector competency in our study was measured through infection rate and *C*Las titer for each individual in each population studied. Studies have shown that approximation of *C*Las titer in individual insects can predict vector capacity. There is a strong positive correlation between *C*Las titer in the psyllid and transmission efficiency (Ammar et al. 2018; Hosseinzadeh et al. 2019a). Broadly, our study shows that there are population-level differences in *D. citri* vector competency that are likely maintained through a combination of gene by environment interactions.

During circulative propagative transmission, *C*Las must cross the gut epithelial cell layer to enter the body cavity of the psyllid and to be considered “acquired”. Next, *C*Las must circulate throughout the body and replicate in tissues, especially the salivary glands, where it has been shown to reach high relative titers. From the salivary glands, transmission is possible and can be inoculated into healthy citrus plants while feeding. The efficiency of the first and second steps can be approximated by using qPCR to detect *C*Las titer in adults that have acquired *C*Las, while the third step can only be measured by careful transmission studies. The development of vector-competent adults requires *C*Las acquisition to occur during the psyllid’s nymphal stages (Inoue et al. 2009; Ammar et al. 2016; Igwe et al. 2022). This study approximates the grove-wide vector capacity of four *D. citri* populations through measurements of *C*Las titer in individual adult insects. A key assumption of this study is that the majority of trees in all citrus groves of Florida are infected with *C*Las and thus, all adult psyllids will have fed on *C*Las-infected citrus as nymphs and are competent transmitters as adults. However, even if this assumption did not hold for every individual in our dataset, multiple studies on vector acquisition rates in the lab have recorded <100% acquisition, despite feeding on known infected material (Pelz-Stelinski et al. 2010; De Souza Pacheco et al. 2020). Samples that do test positive for *C*Las are competent transmitters and fall into categories of low to high vector capacity, as determined by their mean *C*Las titer (Ammar et al. 2018).

Many factors may influence *D. citri* vector competency and the variation we observed in our study. Prior work has shown that host switching influences variation in *D. citri* vector competency (Ramsey et al. 2022; Lombardi et al. 2024). While grove-collected sampling makes it difficult to control or understand all variables that may influence *C*Las titer in psyllids, all four *D. citri* populations were sampled from the same citrus variety (*Citrus sinensis*), within the same week in October, 2018, so it represents a snapshot in time of the potential of *D. citri* in these regions to acquire and transmit. Additionally, the role that symbionts like wDi might play in mediating vector-pathogen interactions is unclear, so the effect of wDi on *C*Las titer from these four fields was also tested. Our results show that *D. citri* has higher wDi titer in CLas-negative individuals, while field site has the strongest effect on *C*Las titer, and sex does not have any effect. The field effect could be related to the heritability of *D. citri* genes involved in vector competency, the management regimen, or transient access to and feeding on non-citrus host plants such as *Murraya spp.* in the grove agro-ecosystem. More work is required to fully explore these hypotheses.

Abandoned groves have also been shown to maintain high *D. citri* populations that travel frequently between managed and abandoned areas (Sétamou et al. 2022; Boina et al. 2009) with equally high *C*Las titers and infection rates as alive and managed groves (Hall 2018). Prior to 2010, Tiwari et al. (Tiwari et al. 2010) studied the *D. citri-C*Las infection rate and found 1.9 % in abandoned and 1.2 % in managed grove populations while the mean *C*Las Cq value was 32.52 in abandoned and 25.95 in managed groves. Over 10 years after Tiwari et al. collected their data, the infection rates found by our study in the least infected, but still managed, grove are at least 1.5x higher incidence while our other three groves have at least 50x higher infection rates. This is undoubtedly due to the vastly increased number of both managed and abandoned infected trees in Florida over the past ten years. A study by De Souza Pacheco et al. (De Souza Pacheco et al. 2020) shows that when *D. citri* are raised on *C*Las-infected citrus for four consecutive generations in a greenhouse setting, the percent-infection and the titer in individuals increased with each generation from 42% to 100% infection incidence, and 47.9 to 5,390 *C*Las cells per 100 ng of DNA. These results suggest that interruptions in vector generations (such as through regular insecticide applications or during seasons of low flush) in the field may help keep *C*Las titer and incidence lower in vector populations. The increase in infection incidence seen in Florida groves despite various management innovations over the last decade is a testament to both the effect of continuous populations of *D. citri* replicating on infected citrus and reference to the accumulating impact HLB has wrought on the Florida citrus industry.

Vector competency of *D. citri* populations and individuals varies widely in the field as we have seen through this study and others (Capoor et al. 1974; Coy and Stelinski 2015; Hall 2018; Ukuda-Hosokawa et al. 2015; Ammar et al. 2018; Sétamou et al. 2016; Xu et al. 1988), in addition to laboratory-kept colonies (Inoue et al. 2009; Hosseinzadeh et al. 2019b; De Souza Pacheco et al. 2020). In agreement with our results, Chu et al (Chu et al. 2016) show that three wild populations of *D. citri* (including a total of 123 insects collected in 2015) vary in their *C*Las titer and *C*Las infection incidence, and that like our study, population (grove site) has the strongest effect on all their models. Coy and Stelinski collected their samples in 2014 from six locations and showed large vector capacity variation ranging from 35-100%, but they collected less than 300 samples in total (Coy and Stelinski 2015). Hall published results from two studies with multiple time points and similar numbers of samples as the current study, but only from one location, and again showed strong variability (Hall 2018). This study not only adds to the literature on in-field vector variability, but it fills a gap in the literature by studying multiple locations with over 150 samples from each location, in 2018 – the most recent timepoint for a study like this, available to our knowledge in the literature. One study focusing on overwintering survival rates assessed psyllids in northern Florida from 2017-2018, but their measurements of *C*Las titer were based on sticky card captures and did not record any psyllids from commercial groves (Martini et al. 2020). Finally, a recent paper, published in 2023 by Ebert et al., surveyed multiple commercial sites in Florida. However, their samples were collected between 2008 and 2012, (Ebert et al. 2023) making our study the latest published record of commercial psyllid vector capacity variability in Florida and perhaps the last one before the wide-spread adoption of injectable oxytetracycline (OTC) for HLB management in Florida. Now that direct injection of the OTC is becoming a wide-spread HLB management tactic in Florida, understanding the impact of OTC on *C*Las titer in psyllids will be important to understand if the use of OTC will indirectly lower vector competency in psyllid populations.

While grove is highly correlated in the current study with *C*Las titer, the base differences may still come down to molecular or genetic differences between the psyllid populations, *C*Las strains, other endosymbionts, and unknown environmental factors. The ability of *D. citri* to acquire and transmit *C*Las may be genetically linked (Ammar et al. 2018) and studies that separate acquisition and transmission (Ammar et al. 2016; Ukuda-Hosokawa et al. 2015; Ammar et al. 2018; Xu et al. 1988) agree that these are likely two separate traits, but that they are correlated, such that higher acquisition rates often lead to more efficient transmission (Ammar et al. 2018). The current study used *C*Las titer as a measure of acquisition in adult *D. citri*, (which may also include effects of internal *C*Las replication) but the high numbers of infected individuals (infection rate) also suggests transmission of *C*Las is occurring in the field and that adult psyllids acquired as nymphs from infected trees.

The relationship between wDi titer on *C*Las in *D. citri* requires further investigation. However, in a related psyllid, *Cacopsylla pyri* (pear psyllid) studied by Serbina et al (Serbina et al. 2022), wDi titer was reduced in males. Several lab studies investigated interactions among wDi, *D. citri* and *C*Las. Hosseinzadeh et al. (Hosseinzadeh et al. 2019b) found that wDi titer varied between sexes, and *D. citri* had higher titer in males that were infected with *C*Las than non-infected males, though wDi did not significantly differ between *C*Las infected and non-infected populations as a whole. Others found that wDi titer was significantly higher in *C*Las-negative *D. citri*, except when controlling for population effect (Chu et al. 2019; Ren et al. 2018; Chu et al. 2016). Fagen et al (Fagen 2012) showed the opposite trend, that *C*Las has a negative relationship with other *D. citri* obligate endosymbionts, but a positive relationship to wDi regarding titer. Our results, using insects collected from citrus groves instead of greenhouse and chamber-raised populations, contradict some of these previous lab-based results. We saw 100% wDi infection rate among our individuals tested, and the titer of wDi tended to be higher in *C*Las non-infected *D. citri* individuals. These contradictions underscore the importance to not over-generalize the results from lab studies and the importance of field-based inquiries on vector-microbe interactions.

## Conclusion

This snapshot assessment of vector populations showed grove-specific differences in *C*Las infection. Specifically, infection rate and titer of *C*Las within individual *D. citri* varied by grove site, exhibiting three patterns of vector competency. However, no conclusions could be drawn on the components of grove site that influenced *C*Las titer in *D. citri.* Additionally, while there was no impact of sex on *C*Las titer, wDi titers were higher in non-infected individuals. These results contradict some studies using lab-reared psyllid colonies. Unknown environmental factors may contribute to titer levels in insects in the grove and we stress that caution is necessary when applying and interpreting results from lab-reared insects to field-based studies and applications. The data suggest that future studies should prioritize understanding the effect of grove management practices, including the potential benefits of using non-citrus plants that are hosts of *D. citri* or of its natural enemies in the grove. Additionally, understanding the contributions that both environment and psyllid genes – the G x E interaction – have on vector competency will help make all future genetic, genomic and field-based studies of HLB more informed when developing control strategies or other programs aimed at blocking *C*Las transmission by *D. citri*.

## Acknowledgements

We thank Tim Eyrich and Shannon Leahy (Southern Gardens), Dr. David Hall and Kathy Moulton (USDA Agricultural Research Service, retired), for their help, organization, and outreach to the growers which enabled us to collect live psyllids from grove sites in Florida, USA.

## Financial Support

Funding for this research was graciously provided by NIFA-AFRI Predoctoral Fellowship award # 2021-67011-35143 for Marina Mann, USDA ARS CRIS project 8062-22410-007-000-D, and USDA NIFA grants 2020-70029-33176 and 2022-70029-38503.

## Competing Interests

The authors declare that they have no competing interests.

## Notes

### Competing Interest Statement

The authors have declared no competing interest.

### Summary of Updates

Updated Figure 2 based on feedback from reviewers.

## References

Ammar, E.-D., Hall, D. G., Hosseinzadeh, S., and Heck, M. 2018. The quest for a non-vector psyllid: Natural variation in acquisition and transmission of the huanglongbing pathogen ‘Candidatus Liberibacter asiaticus’ by Asian citrus psyllid isofemale lines. PLoS ONE. 13:e0195804.

Ammar, E.-D., Ramos, J. E., Hall, D. G., Dawson, W. O., and Shatters, R. G. 2016. Acquisition, Replication and Inoculation of Candidatus Liberibacter asiaticus following Various Acquisition Periods on Huanglongbing-Infected Citrus by Nymphs and Adults of the Asian Citrus Psyllid. PLoS ONE. 11:e0159594.

Boina, D. R., Meyer, W. L., Onagbola, E. O., and Stelinski, L. L. 2009. Quantifying Dispersal of Diaphorina citri (Hemiptera: Psyllidae) by Immunomarking and Potential Impact of Unmanaged Groves on Commercial Citrus Management. Environ Entomol. 38:1250–1258.

Bové, J. M., Garnier, M., Ahlawat, Y. S., Chakraborty, N. K., and Varma, A. 1993. Detection of the Asian Strains of the Greening BLO by DNA-DNA Hybridization in Indian Orchard Trees and Malaysian Diaphorina citri Psyllids. International Organization of Citrus Virologists Conference Proceedings (1957–2010). 12 Available at: https://escholarship.org/uc/item/0ng306x8 [Accessed July 19, 2023].

Capoor, S. P., Rao, D. G., and Viswanath, S. M. 1974. Greening Disease of Citrus in the Deccan Trap Country and its Relationship with the Vector, Diaphorina citri Kuwayama. International Organization of Citrus Virologists Conference Proceedings (1957–2010). 6 Available at: https://escholarship.org/uc/item/6rm6x1tw [Accessed July 19, 2023].

Chen, Q., Li, Z., Liu, S., Chi, Y., Jia, D., and Wei, T. 2022. Infection and distribution of Candidatus Liberibacter asiaticus in citrus plants and psyllid vectors at the cellular level. Microbial Biotechnology. 15:1221–1234.

Chin, E. L., Ramsey, J. S., Mishchuk, D. O., Saha, S., Foster, E., Chavez, J. D., et al. 2020. Longitudinal Transcriptomic, Proteomic, and Metabolomic Analyses of Citrus sinensis (L.) Osbeck Graft-Inoculated with “ Candidatus Liberibacter asiaticus.” J. Proteome Res. 19:719–732.

Chu, C., Hoffmann, M., Braswell, W. E., and Pelz-Stelinski, K. S. 2019. Genetic variation and potential coinfection of Wolbachia among widespread Asian citrus psyllid (Diaphorina citri Kuwayama) populations. Insect Science. 26:671–682.

Chu, C.-C., Gill, T. A., Hoffmann, M., and Pelz-Stelinski, K. S. 2016. Inter-Population Variability of Endosymbiont Densities in the Asian Citrus Psyllid (Diaphorina citri Kuwayama). Microb Ecol. 71:999– 1007.

Coletta-Filho, H. D., Daugherty, M. P., Ferreira, C., and Lopes, J. R. S. 2014. Temporal Progression of ‘Candidatus Liberibacter asiaticus’ Infection in Citrus and Acquisition Efficiency by Diaphorina citri. Phytopathology®. 104:416–421.

Coy, M. R., and Stelinski, L. L. 2015. Great Variability in the Infection Rate of ‘ Candidatus Liberibacter Asiaticus’ in Field Populations of Diaphorina citri (Hemiptera: Liviidae) in Florida. Florida Entomologist. 98:356–357.

De Souza Pacheco, I., Manzano Galdeano, D., Spotti Lopes, J. R., and Machado, M. A. 2020. Development on Infected Citrus over Generations Increases Vector Infection by ‘Candidatus Liberibacter Asiaticus in Diaphorina citri.’ Insects. 11:469.

Ebert, T. A., Shawer, D., Brlansky, R. H., and Rogers, M. E. 2023. Seasonal Patterns in the Frequency of Candidatus Liberibacter Asiaticus in Populations of Diaphorina citri (Hemiptera: Psyllidae) in Florida. Insects. 14:756.

Fagen, J. R. 2012. Characterization of the Relative Abundance of the Citrus Pathogen Ca. Liberibacter Asiaticus in the Microbiome of Its Insect Vector, Diaphorina citri, using High Throughput 16S rRNA Sequencing. TOMICROJ. 6:29–33.

Garnier, M., and Bove, J. 1983. Transmission of the organism associatied with Citrus Greening disease from sweet orange to periwinkle by dodder. Phytopathology. 73:1358–1363.

Ghosh, D., Kokane, S., Savita, B. K., Kumar, P., Sharma, A. K., Ozcan, A., et al. 2023. Huanglongbing Pandemic: Current Challenges and Emerging Management Strategies. Plants. 12:160.

Guidolin, A. S., and Cônsoli, F. L. 2013. Molecular Characterization of Wolbachia Strains Associated with the Invasive Asian Citrus Psyllid Diaphorina citri in Brazil. Microb Ecol. 65:475–486.

Hall, D. G. 2018. Incidence of “ Candidatus Liberibacter asiaticus” in a Florida population of Asian citrus psyllid. J Appl Entomol. 142:97–103.

Higgins, S. A., Mann, M., and Heck, M. 2022. Strain Tracking of ‘ Candidatus Liberibacter asiaticus’, the Citrus Greening Pathogen, by High-Resolution Microbiome Analysis of Asian Citrus Psyllids. Phytopathology®. 112:2273–2287.

Hoffmann, M., Coy, M. R., Gibbard, H. N. K., and Pelz-Stelinski, K. S. 2014. Wolbachia Infection Density in Populations of the Asian Citrus Psyllid (Hemiptera: Liviidae). Environ Entomol. 43:1215–1222.

Hosseinzadeh, S., and Heck, M. 2023. Variations on a theme: factors regulating interaction between Diaphorina citri and “Candidatus Liberibacter asiaticus” vector and pathogen of citrus huanglongbing. Current Opinion in Insect Science. 56:101025.

Hosseinzadeh, S., Ramsey, J., Mann, M., Bennett, L., Hunter, W. B., Shams-Bakhsh, M., et al. 2019a. Color morphology of Diaphorina citri influences interactions with its bacterial endosymbionts and ‘Candidatus Liberibacter asiaticus’ ed. Janice L. Bossart. PLoS ONE. 14:e0216599.

Hosseinzadeh, S., Shams-Bakhsh, M., Mann, M., Fattah-Hosseini, S., Bagheri, A., Mehrabadi, M., et al. 2019b. Distribution and Variation of Bacterial Endosymbiont and “Candidatus Liberibacter asiaticus” Titer in the Huanglongbing Insect Vector, Diaphorina citri Kuwayama. Microb Ecol. 78:206–222.

Igwe, D. O., Higgins, S. A., and Heck, M. 2022. An Excised Leaf Assay to Measure Acquisition of ‘ Candidatus Liberibacter asiaticus’ by Psyllids Associated with Citrus Huanglongbing Disease. Phytopathology®. 112:69–75.

Inoue, H., Ohnishi, J., Ito, T., Tomimura, K., Miyata, S., Iwanami, T., et al. 2009. Enhanced proliferation and efficient transmission of Candidatus Liberibacter asiaticus by adult Diaphorina citri after acquisition feeding in the nymphal stage. Annals of Applied Biology. 155:29–36.

Jain, M., Fleites, L. A., and Gabriel, D. W. 2017. A Small Wolbachia Protein Directly Represses Phage Lytic Cycle Genes in “Candidatus Liberibacter asiaticus” within Psyllids ed. Katherine McMahon. mSphere. 2:e00171–17.

Kassambara, A. 2020. ggpubr: “ggplot2” based publication ready plots. (R package version 0.4.0). Available at: https://CRAN.R-project.org/package=ggpubr.

Kruse, A., Fattah-Hosseini, S., Saha, S., Johnson, R., Warwick, E., Sturgeon, K., et al. 2017. Combining ’omics and microscopy to visualize interactions between the Asian citrus psyllid vector and the Huanglongbing pathogen Candidatus Liberibacter asiaticus in the insect gut ed. PLoS ONE. 12:e0179531.

Kruse, A., Ramsey, J. S., Johnson, R., Hall, D. G., MacCoss, M. J., and Heck, M. 2018. Candidatus Liberibacter asiaticus Minimally Alters Expression of Immunity and Metabolism Proteins in Hemolymph of Diaphorina citri, the Insect Vector of Huanglongbing. J. Proteome Res. 17:2995–3011.

Kuznetsova, A., Brockhoff, P. B., and Christensen, R. H. B. 2017. lmerTest Package: Tests in Linear Mixed Effects Models. J. Stat. Soft. 82 Available at: http://www.jstatsoft.org/v82/i13/ [Accessed August 24, 2023].

Lombardi, R. L., Ramsey, J. S., Mahoney, J. E., MacCoss, M. J., Heck, M. L., and Slupsky, C. M. 2024. Longitudinal Transcriptomic, Proteomic, and Metabolomic Response of Citrus sinensis to Diaphorina citri Inoculation of Candidatus Liberibacter asiaticus. J. Proteome Res. :acs.jproteome.3c00485.

Manjunath, K. L., Halbert, S. E., Ramadugu, C., Webb, S., and Lee, R. F. 2008. Detection of ‘ Candidatus Liberibacter asiaticus’ in Diaphorina citri and Its Importance in the Management of Citrus Huanglongbing in Florida. Phytopathology®. 98:387–396.

Mann, M., Fattah-Hosseini, S., Ammar, E.-D., Stange, R., Warrick, E., Sturgeon, K., et al. 2018. Diaphorina citri Nymphs Are Resistant to Morphological Changes Induced by “Candidatus Liberibacter asiaticus” in Midgut Epithelial Cells ed. Marvin Whiteley. Infect Immun. 86:e00889–17.

Martini, X., Malfa, K., Stelinski, L. L., Iriarte, F. B., and Paret, M. L. 2020. Distribution, Phenology, and Overwintering Survival of Asian Citrus Psyllid (Hemiptera: Liviidae), in Urban and Grove Habitats in North Florida ed. Blake Bextine. Journal of Economic Entomology. 113:1080–1087.

Naeem, A., Afzal, M. B. S., Freed, S., Hafeez, F., Zaka, S. M., Ali, Q., et al. 2019. First report of thiamethoxam resistance selection, cross resistance to various insecticides and realized heritability in Asian citrus psyllid Diaphorina citri from Pakistan. Crop Protection. 121:11–17.

Nakabachi, A., Ueoka, R., Oshima, K., Teta, R., Mangoni, A., Gurgui, M., et al. 2013. Defensive Bacteriome Symbiont with a Drastically Reduced Genome. Current Biology. 23:1478–1484.

Neupane, S., Bonilla, S. I., Manalo, A. M., and Pelz-Stelinski, K. S. 2022. Complete de novo assembly of Wolbachia endosymbiont of Diaphorina citri Kuwayama (Hemiptera: Liviidae) using long-read genome sequencing. Sci Rep. 12:125.

Pardo, S., Martínez, A. M., Figueroa, J. I., Chavarrieta, J. M., Viñuela, E., Rebollar-Alviter, Á., et al. 2018. Insecticide resistance of adults and nymphs of Asian citrus psyllid populations from Apatzingán Valley, Mexico: Resistance of D. citri to bifenthrin, malathion and chlorpyrifos. Pest. Manag. Sci. 74:135–140.

Pelz-Stelinski, K. S., Brlansky, R. H., Ebert, T. A., and Rogers, M. E. 2010. Transmission Parameters for Candidatus Liberibacter asiaticus by Asian Citrus Psyllid (Hemiptera: Psyllidae). J. Econ. Entom. 103:1531–1541.

R Core Team. 2022. R: A language and environment for statistical computing. Available at: https://www-R-project.org/.

Ramsey, J. S., Ammar, E.-D., Mahoney, J. E., Rivera, K., Johnson, R., Igwe, D. O., et al. 2022. Host Plant Adaptation Drives Changes in Diaphorina citri Proteome Regulation, Proteoform Expression, and Transmission of ‘Candidatus Liberibacter asiaticus’, the Citrus Greening Pathogen. Phytopathology®. 112:101–115.

Ramsey, J. S., Johnson, R. S., Hoki, J. S., Kruse, A., Mahoney, J., Hilf, M. E., et al. 2015. Metabolic Interplay between the Asian Citrus Psyllid and Its Profftella Symbiont: An Achilles’ Heel of the Citrus Greening Insect Vector ed. Murad Ghanim. PLoS ONE. 10:e0140826.

Rao, C. N., Shivankar, V. J., Sandnya, D., David, K. J., and Dhengre, V. N. 2014. Insecticide resistance in field populations of Asian Citrus psyllid, Diaphorina citri Kuwayama (Hemiptera: Psyllidae). Pesticide Research Journal. 26:42–47.

Razi, M. F., Keremane, M. L., Ramadugu, C., Roose, M., Khan, I. A., and Lee, R. F. 2014. Detection of Citrus Huanglongbing-Associated ‘ Candidatus Liberibacter asiaticus’ in Citrus and Diaphorina citri in Pakistan, Seasonal Variability, and Implications for Disease Management. Phytopathology®. 104:257– 268.

Ren, S.-L., Li, Y.-H., Ou, D., Guo, Y.-J., Qureshi, J. A., Stansly, P. A., et al. 2018. Localization and dynamics of Wolbachia infection in Asian citrus psyllid Diaphorina citri, the insect vector of the causal pathogens of Huanglongbing. MicrobiologyOpen. 7:e00561.

Saha, S., Hunter, W. B., Reese, J., Morgan, J. K., Marutani-Hert, M., Huang, H., et al. 2012. Survey of Endosymbionts in the Diaphorina citri Metagenome and Assembly of a Wolbachia wDi Draft Genome ed. Dan Zilberstein. PLoS ONE. 7:e50067.

Serbina, Š. L., Gajski, D., Malenovský, I., Corretto, E., Schuler, H., and Dittmer, J. 2022. Wolbachia infection dynamics in a natural population of the pear psyllid Cacopsylla pyri (Hemiptera: Psylloidea) across its seasonal generations. Sci Rep. 12:16502.

Sétamou, M., Alabi, O. J., Kunta, M., Dale, J., and Da Graça, J. V. 2020. Distribution of Candidatus Liberibacter asiaticus in Citrus and the Asian Citrus Psyllid in Texas Over a Decade. Plant Disease. 104:1118–1126.

Sétamou, M., Alabi, O. J., Kunta, M., Jifon, J. L., and Da Graça, J. V. 2016. Enhanced Acquisition Rates of ‘ Candidatus Liberibacter asiaticus’ by the Asian Citrus Psyllid (Hemiptera: Liviidae) in the Presence of Vegetative Flush Growth in Citrus. J Econ Entomol. 109:1973–1978.

Sétamou, M., Tarshis Moreno, A., and Patt, J. M. 2022. Source or Sink? The Role of Residential Host Plants in Asian Citrus Psyllid Infestation of Commercial Citrus Groves ed. Arash Rashed. Journal of Economic Entomology. 115:438–445.

Shippy, T. D., Hosmani, P. S., Flores-Gonzalez, M., Mann, M., Miller, S., Weirauch, M. T., et al. 2019. Diaci v3.0: Chromosome-level assembly, de novo transcriptome and manual annotation of Diaphorina citri, insect vector of Huanglongbing. Genomics. Available at: http://biorxiv.org/lookup/doi/10.1101/869685 [Accessed August 24, 2023].

Tiwari, S., Lewis-Rosenblum, H., Pelz-Stelinski, K., and Stelinski, L. L. 2010. Incidence of Candidatus Liberibacter asiaticus Infection in Abandoned Citrus Occurring in Proximity to Commercially Managed Groves. jnl. econ. entom. 103:1972–1978.

Ukuda-Hosokawa, R., Sadoyama, Y., Kishaba, M., Kuriwada, T., Anbutsu, H., and Fukatsu, T. 2015. Infection Density Dynamics of the Citrus Greening Bacterium “Candidatus Liberibacter asiaticus” in Field Populations of the Psyllid Diaphorina citri and Its Relevance to the Efficiency of Pathogen Transmission to Citrus Plants ed. H. Goodrich-Blair. Appl Environ Microbiol. 81:3728–3736.

Vázquez-García, M., Velázquez-Monreal, J., Medina-Urrutia, V. M., Cruz-Vargas, C. D. J., Sandoval-Salazar, M., Virgen-Calleros, G., et al. 2013. Insecticide Resistance in Adult Diaphorina citri Kuwayama from Lime Orchards in Central West Mexico. Southwestern Entomologist. 38:579–596.

Venkatesan, T., Chethan, B. R., and Mani, M. 2022. Insecticide Resistance and Its Management in the Insect Pests of Horticultural Crops. In Trends in Horticultural Entomology, ed. M. Mani. Singapore: Springer Nature Singapore, p. 455–490. Available at: https://link.springer.com/10.1007/978-981-19-0343-4_14 [Accessed August 24, 2023].

Walter, A. J., Hall, D. G., and Duan, Y. P. 2012. Low Incidence of ‘ Candidatus Liberibacter asiaticus’ in Murraya paniculata and Associated Diaphorina citri. Plant Disease. 96:827–832.

Wickham, H. 2-16. ggplot2: Elegant graphics for data analysis.

Wulff, N., Daniel, B., Sassi, R., Moreira, A., Bassanezi, R., Sala, I., et al. 2020. Incidence of Diaphorina citri Carrying Candidatus Liberibacter asiaticus in Brazil’s Citrus Belt. Insects. 11:672.

Xu, C. F., Xia, Y. H., Li, K. B., and Ke, C. 1988. Further Study of the Transmission of Citrus Huanglungbin by a Psyllid, Diaphorina citri Kuwayama. International Organization of Citrus Virologists Conference Proceedings (1957-2010). 10 Available at: https://escholarship.org/uc/item/0w42q0r7 [Accessed July 19, 2023].

